# Single-Cell Analysis of the 3D Topologies of Genomic Loci Using Genome Architecture Mapping

**DOI:** 10.1101/2020.02.10.941047

**Authors:** Lonnie R. Welch, Catherine Baugher, Yingnan Zhang, Trenton Davis, William F. Marzluff, Joshua D. Welch, Ana Pombo

**Affiliations:** School of Computer Science and Electrical Engineering, Ohio University, Athens, Ohio, USA; Dept. of Biochemistry and Biophysics and Integrative Pgm. in Biological and Genome Sciences, University of North Carolina, Chapel Hill, NC 27599; Department of Computational Medicine and Bioinformatics, University of Michigan, Ann Arbor, Michigan, USA; Berlin Institute for Medical Systems Biology, Max Delbrück Center for Molecular Medicine and Humboldt University of Berlin, Berlin, Germany

## Abstract

Although each cell within an organism contains a nearly identical genome sequence, the three-dimensional (3D) packing of the genome varies among individual cells, influencing cell-type-specific gene expression. Genome Architecture Mapping (GAM) is the first genome-wide experimental method for capturing 3D proximities between any number of genomic loci without ligation. GAM overcomes several limitations of 3C-based methods by sequencing DNA from a large collection of thin sections sliced from individual nuclei. The GAM technique measures locus co-segregation, extracts radial positions, infers chromatin compaction, requires small numbers of cells, does not depend on ligation, and provides rich single-cell information. However, previous analyses of GAM data focused exclusively on population averages, neglecting the variation in 3D topology among individual cells.

We present the first single-cell analysis of GAM data, demonstrating that the slices from individual cells reveal intercellular heterogeneity in chromosome conformation. By simultaneously clustering both slices and genomic loci, we identify topological variation among single cells, including differential compaction of cell cycle genes. We also develop a geometric model of the nucleus, allowing prediction of the 3D positions of each slice. Using GAM data from mouse embryonic stem cells, we make new discoveries about the structure of the major mammalian histone gene locus, which is incorporated into the Histone Locus Body (HLB), including structural fluctuations and putative causal molecular mechanisms. Our methods are packaged as *SluiceBox*, a toolkit for mining GAM data. Our approach represents a new method of investigating variation in 3D genome topology among individual cells across space and time.

## 1. Introduction

The higher-order structure of chromatin can be understood in terms of 3D genomic units that associate with each other in a compartmentalized fashion, producing Topologically Associated Domains (TADs). Recent studies suggest that TADs and chromatin loops are dynamic structures which may coalesce dynamically into various topologies over the course of the cell cycle [1]. Indeed, there are several ways in which the chromatin structure within a single cell may fluctuate. The cell cycle is perhaps the most accessible example of such fluctuation, as previous single-cell Hi-C studies have observed by mapping cell cycle specific structural changes in genome organization [2].

A cell’s chromatin structure may also be impacted by signal transduction, wherein extracellular stimuli can trigger pathways to activate or repress sets of target genes. Changes to chromatin structure may occur through these signaling processes by the targeting of modifications to histone N-terminal tails [3]. This allows cells to dynamically change chromatin structure in order to regulate genes situationally. An example of this mechanism in action can be seen in the targeted activation of genes that initiate the inflammatory response [4]. The gene products induced by signal transduction may be used to directly target foreign substances, or to recruit cells of the immune system to defend a specific region. On a smaller scale, gene expression variation may relate to structural changes. For example, microscopic images of NANOG in pluripotent cells show fluctuations in expression levels, which may correspond to chromatin structural changes.

Although general, cell-wide changes in the 3D architecture of the genome are widely recognized, it is not known how the chromatin structure fluctuates in accordance with cell states, such as the cell cycle. This necessitates reliable single-cell techniques that detect when 3D genome architecture changes and how such variations relate with the physiological state of the cell.

The independent sequencing of individual nuclear slices in Genome Architecture Mapping (GAM) [7] provides the opportunity to explore single-cell behaviors in 3D chromatin folding. GAM was previously used to study mouse embryonic stem cells (mESCs), which are actively dividing pluripotent cells. Through the sequencing of the DNA content of only 408 nuclear profiles (NPs), the previous GAM study [7] revealed new features of chromatin contacts that were not seen by Hi-C, including extensive contacts between enhancers and active genomic regions that span tens of megabases. In this study, we explore the power of GAM for capturing fluctuations of chromatin architecture within a population of cells. We use GAM sequencing data from single-cell nuclear profiles collected from mESCs to investigate variations in genome organization across cells, and the implications they may have on the regulation of gene expression. We consider a region that encodes the mammalian Hist1 cluster, which spans several Mb, contains multiple copies of the genes for the five different histone proteins, and includes several sub-groups of associated histone genes. The coordinate regulation of multiple histone genes during the cell cycle is well-established, and predicts distinct cell cycle-regulated states of histone gene expression. The Hist1 cluster is a relatively small region of the genome, which undergoes dynamic regulation in all cells, that is amenable to study by GAM. Single-cell states of 3D chromatin structure of the histone locus serve as a case study to better understand the mechanisms of transcription of histone genes and how the dynamic 3D architecture of chromatin may affect relatively small-scale genomic regions.

The histone locus body (HLB) assembles at histone genes, and is a nuclear body where the production of histone mRNA occurs and has been observed at two different loci in mammalian cells [5,10,11]. The genes that encode histone proteins occur within four genomic clusters. Hist1 is the largest cluster, encoding representatives of all five classes of histone proteins and containing 55 histone genes in humans and 51 histone genes in mouse. Hist2 contains 10-12 genes for the four core histone proteins. The genes in the Hist1 cluster are arranged into three gene-rich subclusters that span ~2.5 Mb. A large gap of roughly 1.3 Mb exists between two of the subclusters, where many other genes reside but histone genes are absent. A second HLB assembles at the Hist2 cluster which spans about 100 Kb and contains 10-12 genes in mammals. In this study we focus on the Hist1 region. We hypothesize that there are multiple cell states—for example, due to cell cycle stage—which influence the regulation of genes in the Hist1 region, resulting in distinct chromatin configurations. Cell cycle regulation is a vital component of the proper function of nucleosomes. Transcription of core histone genes increases sharply as cells enter S-phase and decreases shortly after S-phase, allowing cells to package newly replicated DNA [6]. This is achieved through changes in the HLB structure, allowing for ample recruitment of transcription factors and proteins related to RNA synthesis during S-phase. When the normal processes of the HLB are impaired, genomic instability may result, putting the cell at risk of oncogenesis.

Although the GAM method inherently performs single-cell chromatin co-segregation sampling, GAM data have not so far been used for single-cell studies of chromatin architecture. Thus, in this paper we present *SluiceBox*, a novel pipeline for studying the genome architecture of selected loci. We chose the name SluiceBox—the water-and-trough system used by miners to pan for gold—because it serves as an accurate metaphor for mining GAM data for nuggets of insight into chromatin conformation. We employ *SluiceBox* to analyze single-cell GAM data in order to study the Hist1 region. We find that the Hist1 region does show states with distinctly different chromatin structures. We use GAM and *SluiceBox* to explore the hypothesis that the various states we observed originate from differences in cell cycle phase.

## 2. Methods

This section describes *SluiceBox*, a software pipeline that analyzes single-cell GAM data to discover chromatin states of specific genomic loci; to characterize the states in terms of nuclear organization, genomics, and epigenetics; and to infer the putative structures, functions and mechanisms pertaining to chromatin interaction. As shown in Figure 1, the input to *SluiceBox* is produced by GAM [7] and GAM-tools [9], which generate a collection of slices from single cells. The major phases of the *SluiceBox* pipeline are: (1) *cluster generation*, (2) *feature generation*, and (3) *interpretation*.

**Figure 1.**
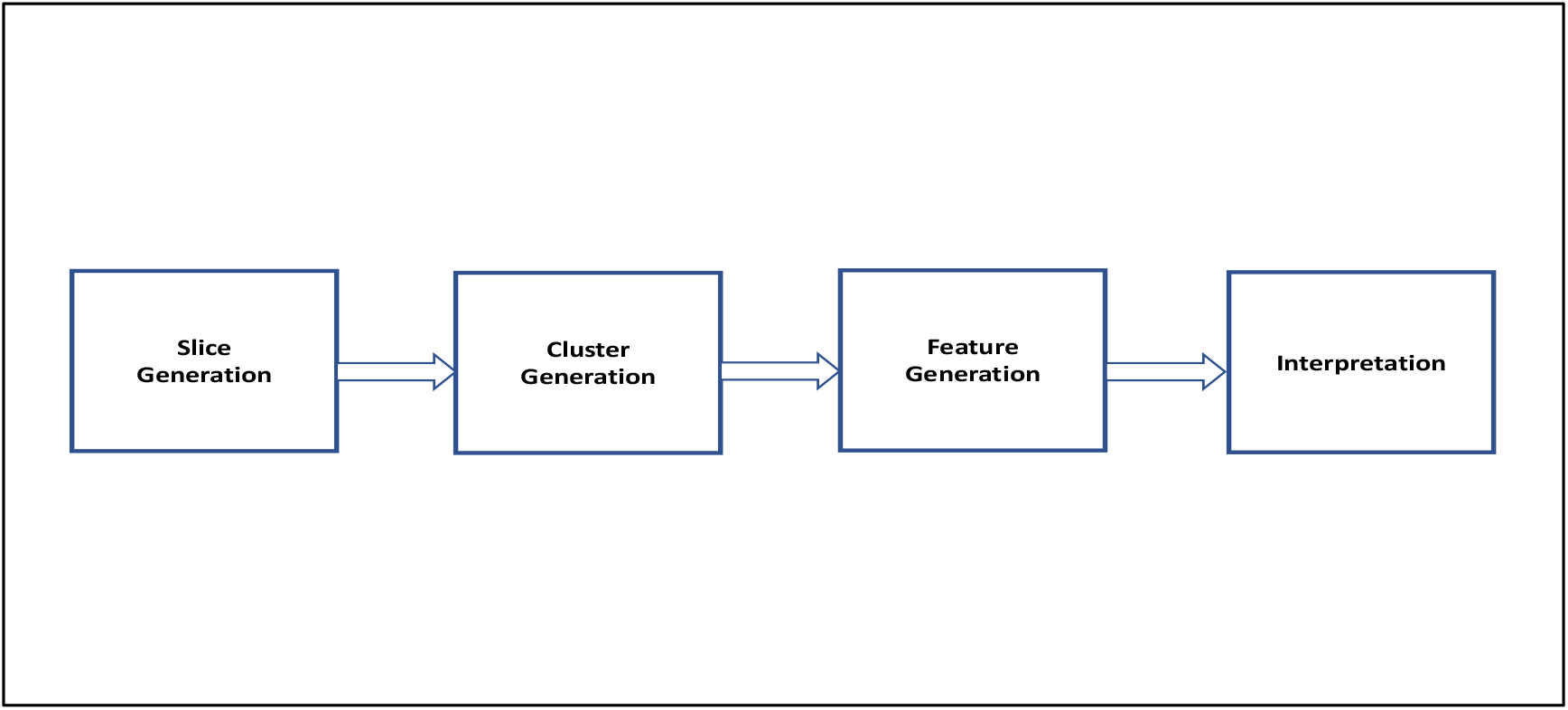
The experimental and computational pipeline for single-cell analysis of the 3D topologies of genomic loci of interest via Genome Architecture Mapping (GAM).

### 2.1 Cluster Generation

GAM-tools produces a segmentation table (**ST**) which represents the genome-wide interactions captured by GAM. Each row in the table represents a specific genomic window and each column corresponds to a nuclear profile (NP), i.e., a slice from a single cell. Each NP originates from a separate cell. Each entry in the table indicates whether a particular genomic window was captured by a specific NP. One can utilize the array of single-cell information produced by GAM to study the chromatin organization within a genomic region of interest (GRI).

Data cleaning is performed to remove windows with very low detection frequency or very high detection frequency. In the case study, windows with detection frequency greater than 100 (out of 480) are removed. Exploratory analysis is performed to ensure that (1) the genomic region to be studied is captured by NPs in **ST** and (2) there is variation among the relevant NPs. Exploration begins by extracting from **ST** all slices that capture the GRI. Let **S** represent the set of selected slices. It is recommended that each selected NP captures a significant portion (e.g., >= 15%) of the region of interest. Assess the number of NPs that intersect the GRI, the number of windows within the GRI that are captured by the NPs in **ST**, and, for each window within the GRI, the number of NPs that detected the window.

To assess variation among NPs for the GRI, a similarity matrix is generated by applying a summed AND operator between pairwise NPs, and a principal component analysis (PCA) is performed by applying Singular Value Decomposition on the segmentation table with two components. To avoid weighting some NPs more heavily than others, the similarity matrix is normalized by dividing each comparison by the lowest total number of windows in a pair of NPs. This matrix is visualized as a heatmap and clustered along both the x-axis and the y-axis using hierarchical clustering with a Euclidean distance metric and single linkage method. The PCA plot and heatmap may be used to perform an initial assessment to determine whether interesting patterns exist in the data, before proceeding with a determination of where different configurations of chromatin contacts are captured.

Assuming that exploratory analysis indicates that interesting patterns may exist within the data, identify each homogeneous subset of slices that captures a unique pattern of contacts within the region of interest. To accomplish this task, partition **S** into subsets {s_1_, s_2_, s_3_, … s_n_}, wherein

- each s_i_ consists of slices that captured similar patterns of contacts within the region of interest and
- the patterns of contacts captured in s_i_ are different from those captured in s_j_ (for all *i*,*j* where *i*<>*j*).

Each subset contains NPs that were clustered together because they captured similar chromatin interaction patterns in the region of interest. K-means clustering is used to cluster NPs that represent putative cell states. The number of clusters to generate (i.e., ‘k’) can be informed by results obtained in the exploratory analysis steps of the pipeline. Different configurations of chromatin contacts can be visualized in a clustered heatmap of NPs, with each cluster representing a distinct set of chromatin contacts. Some pairs of clusters may contain complementary sets of windows, indicating a chromatin configuration found in a subset of the cells, jointly representing a putative cell state. A cluster containing all windows in the GRI may capture a cell state wherein the chromatin is compacted during cell division, or for small genomic regions it may represent the presence of all genomic windows within the physical slice, independently of its configuration. The set of windows associated with a cluster of NPs represents genomic loci that interact in a subset of cells; this set of windows can be visualized as a track in a genome browser. The simultaneous viewing of multiple NP clusters in a genome browser can reveal interesting genomic patterns that represent fluctuations in chromatin conformation – i.e., states.

### 2.2 Feature Generation

To gain clues about the functional characteristics of a cluster of cells, available -omics data (e.g., histone marks, TF binding, chromatin accessibility, and lamina association) are associated with the set of loci captured by a cluster and are visualized in a genome browser. Additionally, 3D position and chromatin compaction are estimated for each cluster, leveraging the ability of GAM to create a rich single-cell model that is beyond what is possible with single-cell Hi-C.

#### 2.2.1 Estimating 3D Position

Because each nuclear profile is generated by slicing an individual nucleus, it is possible in principle to use a geometric model of the nucleus to estimate the position from which each slice was taken. The original GAM paper used a simple model, assuming that the nucleus is a sphere, allowing calculation of the “radial position”, or distance from the sphere’s equator, for each slice. To calculate radial position of an NP, we extended the method described in [7] for single-cell analyses. In brief, the authors demonstrated that the percentage of the genome covered by each NP can be used as a proxy for its latitude relative to the most equatorial NPs on the sphere representing the shape of the nucleus. Thus, equatorial NPs are those that cover the largest portion of the genome and apical NPs are those that cover the smallest portion of the genome. Radial position for each NP is computed based on the number of loci detected:

1. For each NP, compute the number of windows that it detected (across the entire genome).
2. Estimate the radial position of each NP:
  a. Rank the NPs by number of windows detected (across the entire genome).
  b. Consider the top **Θ** percent of ranked NPs to be equatorial, the bottom **Π** percent of ranked NPs to be apical, and all other NPs to be neither equatorial nor apical. The default value for **Θ** and **Π** is 25%.

The radial position of an NP cluster can be estimated by considering the radial positions of the NPs in the cluster. There are several possibilities. Is a cluster made up of NPs from homogeneous radial positions? If so, the cluster captures a specific type of slice - either apical of equatorial slices (or somewhere in between apical and equatorial); this may yield insight regarding the function or state of the cells captured by the NPs. If a cluster consists of NPs from diverse radial positions, then it may be useful to partition it into subclusters based on radial position. Comparing the radial positions of clusters may provide insight into the degree of fluctuation of the locus; further investigation (see section 3.3) may explain the functional implications of the different chromatin configurations represented by the clusters.

We also derived a more general version of the geometric model used in the GAM paper by modeling the nucleus as an ellipsoid rather than a sphere. We represent each slice as the intersection between two parallel planes separated by the slice width, **s** = 0.22*μ*m. The volume of the slice is then determined by where the planes intersect the ellipsoid. The key insight of our approach is that, by calculating the proportion of the ellipsoid’s volume captured by each slice, we can use the equation for the intersection between a plane and an ellipsoid to determine the plane that yields the observed volume. The details of the mathematical derivation for this approach are shown below. Our method computes the volume of intersection between two parallel planes and an ellipsoid, given an ellipsoid 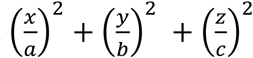 with known parameters *a*,*b*,*c*, and a plane *mx* + *ny* + *kz* = 0. The intersection between the ellipsoid and plane is then an ellipse with equation:

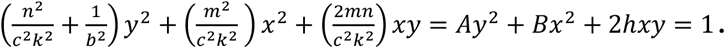

The area of this ellipse is: 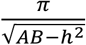

Now if we consider a slice formed by two parallel planes separated by a distance of **s**, we can approximate the volume of the slice with an elliptical cylinder. This neglects the curvature of the ellipsoid between the two planes, but since **s** is small, this approximation should be quite accurate. The volume of the slice is then: 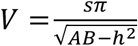. In terms of the original ellipsoid and plane parameters, this equation is:

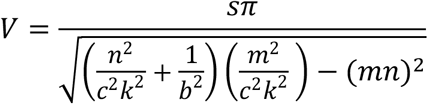

We can estimate the volume of a given slice by calculating the fraction of the genome captured in the slice; this slice then represents the same fraction of the total ellipsoid volume. We can then use the above equation to solve for the equation of the plane that yields the observed volume.

#### 2.2.2 Estimating Chromatin Compaction

To estimate chromatin compaction for single-cell GAM data, we extend the method described in [7], which relies on the following observation:

> “We reasoned that de-condensed genomic loci should occupy larger volumes (or adopt more elongated conformations) than more condensed loci. De-condensed loci would therefore be intersected more frequently (and be detected more frequently in randomly-oriented nuclear profiles) than smaller or more spherical loci.” [7]

This chromatin compaction estimation method does not represent a single-cell perspective – it is for pools of cells. Thus, we present a new approach, which computes compaction for each cluster, using the insight that decondensed loci occupy large volumes and therefore are intersected more frequently than smaller loci.

Procedure for calculating chromatin compaction for a cluster of NPs:

A. for each window *i* detected *within* a cluster **c** (denoted as **w**_**i,c**_),
  1. Compute NP(**w**_**i,c**_) – the number of NPs within cluster **c** that detected window *i*. <u>NOTE</u>: A window *i* is considered to be detected within a cluster **c** if that window occurs in at least **μ**% of the NPs in cluster **c**. The default value for **μ** is 25%. <u>NOTE</u>: To avoid bias due to the sizes of NP clusters (which may cause small clusters to be considered to have high compaction), the value of NP(**w**_**i,c**_) is normalized, by dividing it by the total number of NPs in cluster **c**.
B. for **W_c_**={ **w**_**i,c**_, **w**_**j,c**_, … **w**_**z,c**_}, the set of all windows detected *within* a cluster **c**
  1. display a box plot NP_c_ of the NP(**w**_**j,c**_) values, for all *i*
  2. compute mean(**W**_**c**_) – the mean of NP(**w_i,c_**), for all windows *i* that were detected by NPs in cluster **c**. That is, compute the mean value of this set: {NP(**w**_**i,c**_), NP(**w**_**j,c**_), … NP(**w**_**z,c**_)}
C. Compare the mean(**W**_**c**_) values of all clusters, **c**. Decondensed loci are intersected more frequently (i.e., have larger mean(**W**_**c**_) values) than smaller loci.
  1. compare box plots of NP_c_, for all *c*
  2. generate a box plot of the mean(**W_c_**) values of all clusters, **c**

### 2.3 Interpretation

Given a set of cell clusters that have been annotated with estimated radial position and chromatin compaction, along with genome browser views of (1) the genomic windows in each cluster and (2) -omics data, one can make inferences about the chromatin structure, the function, and the mechanistic drivers of each putative cell state (recall that each cluster of NPs represents a putative *cell state*).

The set of genomic windows captured by a cluster implies a particular chromatin structure (intra-cluster structure) which can be inferred by window interaction analysis. The chromatin structure implied by *interactions within each cluster* is that all blocks of contiguous windows in the cluster are considered to interact with each other. To visualize these interactions, a graph can be deduced, wherein each block is a node and each interaction is an edge.

Interestingly, some pairs of clusters may identify complementary sets of windows in the GRI. The windows of each cluster within such a pair represent a putative chromatin substructure that is formed in a particular cell state; the union of the complementary substructures represents the structure of the GRI in a putative state. In a similar manner as for intra-cluster structures, a complementary set of clusters can be used to perform structural analysis to identify an inter-cluster structure. All complementary clusters are considered to interact with each other, and a graph can be inferred, wherein each block of contiguous genomic windows is a complementary cluster and each inter-cluster interaction is an edge.

Given the inferred chromatin structures, it is useful to consider carefully the -omics features associated with each cluster. To provide insight into the purpose and mechanisms of formation of the inferred structures, cluster membership can be correlated with epigenetic and genomic features, radial position and compaction. We have found it valuable to classify each cluster of NPs using the features. This has yielded hypotheses about the putative functions of a set of cells as well as possible mechanisms for the formation of a specific chromatin configuration. Additional insight can be gained by performing functional analysis for each cluster, identifying the specific genes, gene families, enhancers, and pathways that are present in each cluster and in each complementary set of clusters. This may yield insights into the possible reason for the inferred structure and for the -omics features (e.g., to activate a set of genes that are needed in a particular cell state).

## 3. Results

The GAM single-cell analysis pipeline is used to characterize the Histone Locus Body (HLB), a nuclear body that performs the important role of making histone proteins [5]. The HLB consists of multiple, independent bodies that are formed by chromatin looping within two different regions of the genome (Hist1 and Hist2). During mitosis and cell division the HLB is dissociated, but immediately after cell division the HLB is formed. Only in S-phase are large amounts of histone mRNA produced, with G1 and G2 phases having much lower amounts. In this paper we study the largest of the two HLBs – Hist1. Located on mouse chromosome 13:21.7Mb-24.1Mb (genome build MM9), this GRI was studied using 40Kb resolution GAM nuclear profiles from 480 mouse embryonic stem cells (mESCs) from the 46C line. Our results demonstrate that GAM data can be used to detect fluctuations in the chromatin structure of a locus of interest, and that the *SluiceBox* pipeline provides insights into putative mechanisms that cause chromatin structures to form. Specifically, the results show the power of GAM and the effectiveness of our pipeline by identifying states of Hist1 and characterizing the structure and features of Hist1 in greater detail than has been done previously.

### 3.1. Cluster generation

The Hist1 GRI was extracted from the segmentation table, and NPs that did not capture the GRI were removed. Subsequently, exploratory statistics were computed for NPs, windows, and the GRI; the data were filtered and assessed for degree of variance; and the NPs were clustered. A histogram was initially created to depict the coverage of the GRI by the NPs, providing insight into the distribution of windows across the data and aiding in selection of an appropriate cut-off for filtering NPs. NPs that did not capture at least 10 windows in the Hist1 region were removed, resulting in a set of NPs that cover between 10 windows to at most 53 windows within the GRI (the GRI contains 59 windows of size 40Kb).

A clustered similarity matrix (see Figure 2) shows groups of NPs that result from considering pairwise similarities. The dendrogram shows inter-NP relationships, while the value signifies similarity at each pairwise comparison. In this result, we see groups of NPs clustered when there is high similarity among the NPs, such as the cluster that occurs between NPs F10F1 and F11C2 in the plot. Another group appearing to have a similar property may be seen between NPs listed consecutively from F8C5 through F15E7, and a group lying between F11B1 and F9C5 have strong intra-group similarity and low similarity with other NPs. Furthermore, the PCA (see Figure 3) also clearly shows interesting patterns of variation among NPs, with data points spread out from each other and potentially two or three separate groups appearing to localize together in the figure. Thus, our filtering threshold was not too strict and multiple cell states may, in fact, be captured by the data.

**Figure 2.**
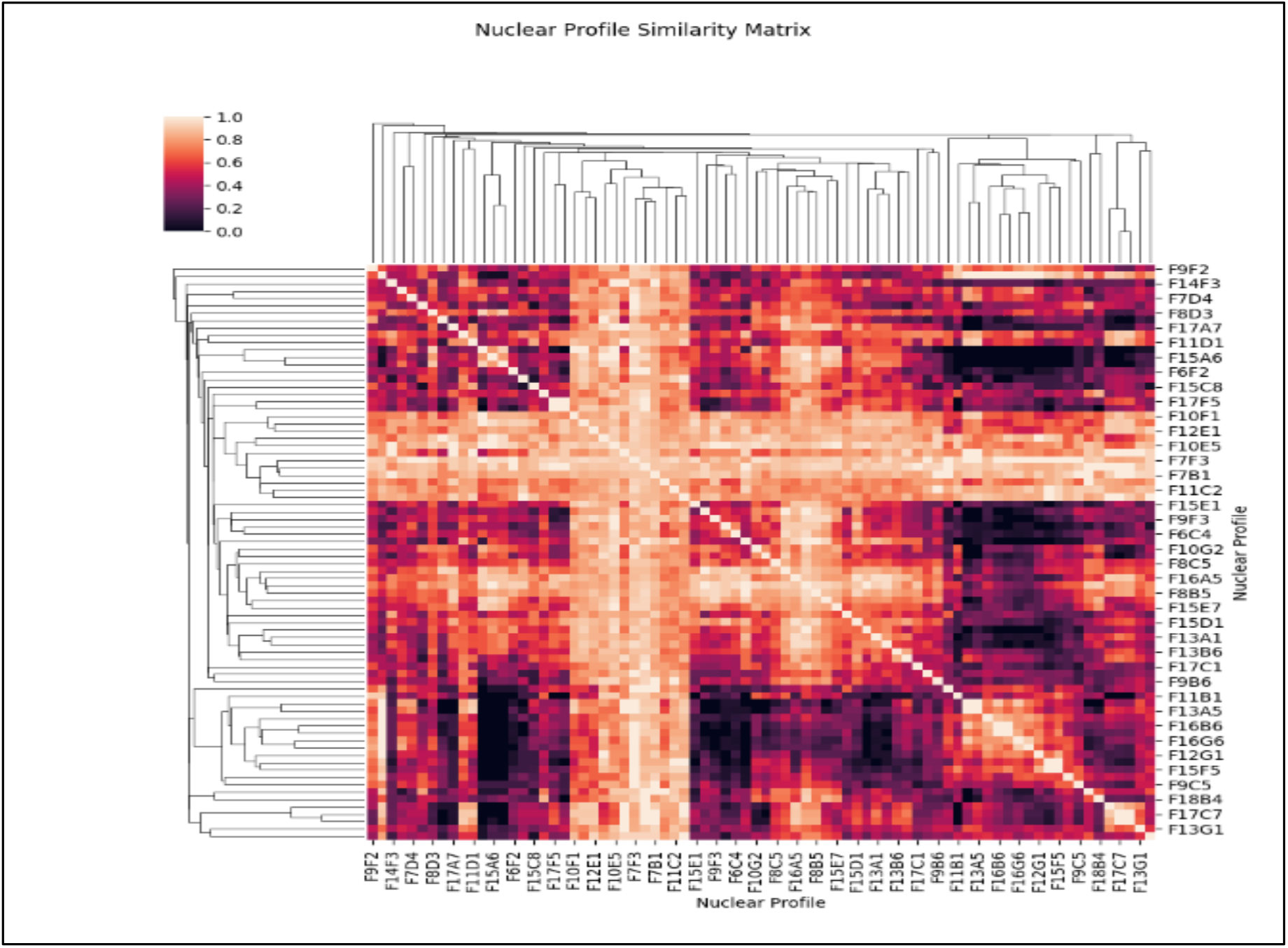
A similarity matrix of NPs that captured the Hist1 region.

**Figure 3.**
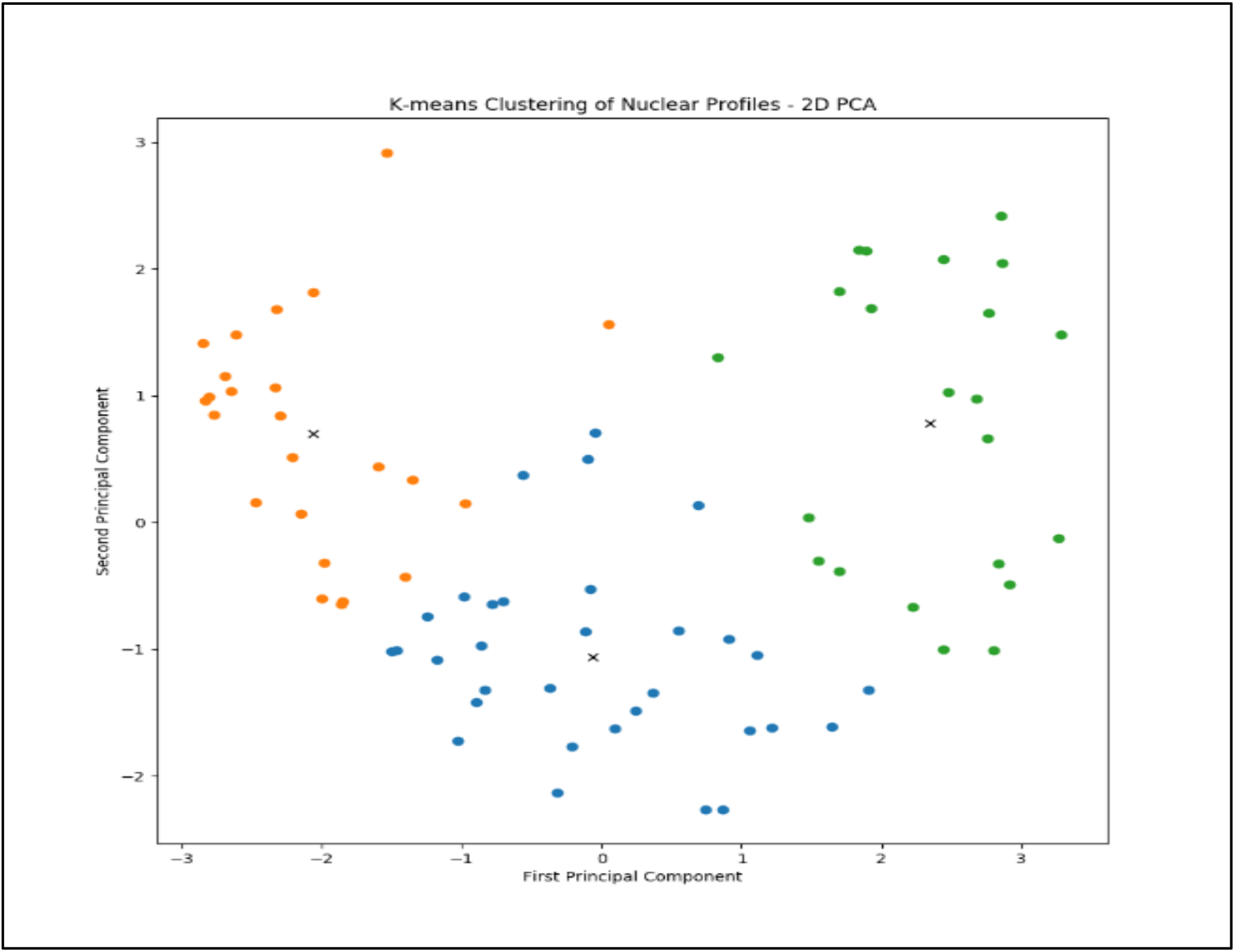
PCA of Nuclear Profiles in Genomic Region of Interest. NPs are colored by cluster membership.

To identify structural variation within the cell population, the NPs were clustered using a k-means clustering algorithm, with a maximum number of 1000 iterations and the *number of clusters* parameter set to 3. The number of clusters parameter was chosen by looking at the PCA and normalized similarity clustered heatmap in the basic statistics step of the pipeline, which showed three groups of NPs. This resulted in subgroups that have different combinations of windows from within the GRI. After performing k-means clustering, labels are obtained for each data point, and NPs are color-coded by cluster. The different configurations of chromatin contacts are reflected in the color-coded PCA of Figure 3. Interestingly, the clusters fall into different regions of the PCA plot, suggesting that the clusters capture strong patterns in the data. The clusters of NPs are shown in the heatmap in Figure 4, wherein one can observe distinct visual patterns of genomic windows in each cluster. A genome browser view of the Hist1 region and the three clusters is shown in Figure 5. Clusters 1 and 2 (depicted as blue and red tracks) span complementary portions of the GRI, and thus they may capture the genome architecture one cell state – denoted as {clust1, clust2}. The green track, which consists of windows that span the entire GRI, is considered one state; this is cluster 3, which may capture the architecture of an alternate state.

**Figure 4.**
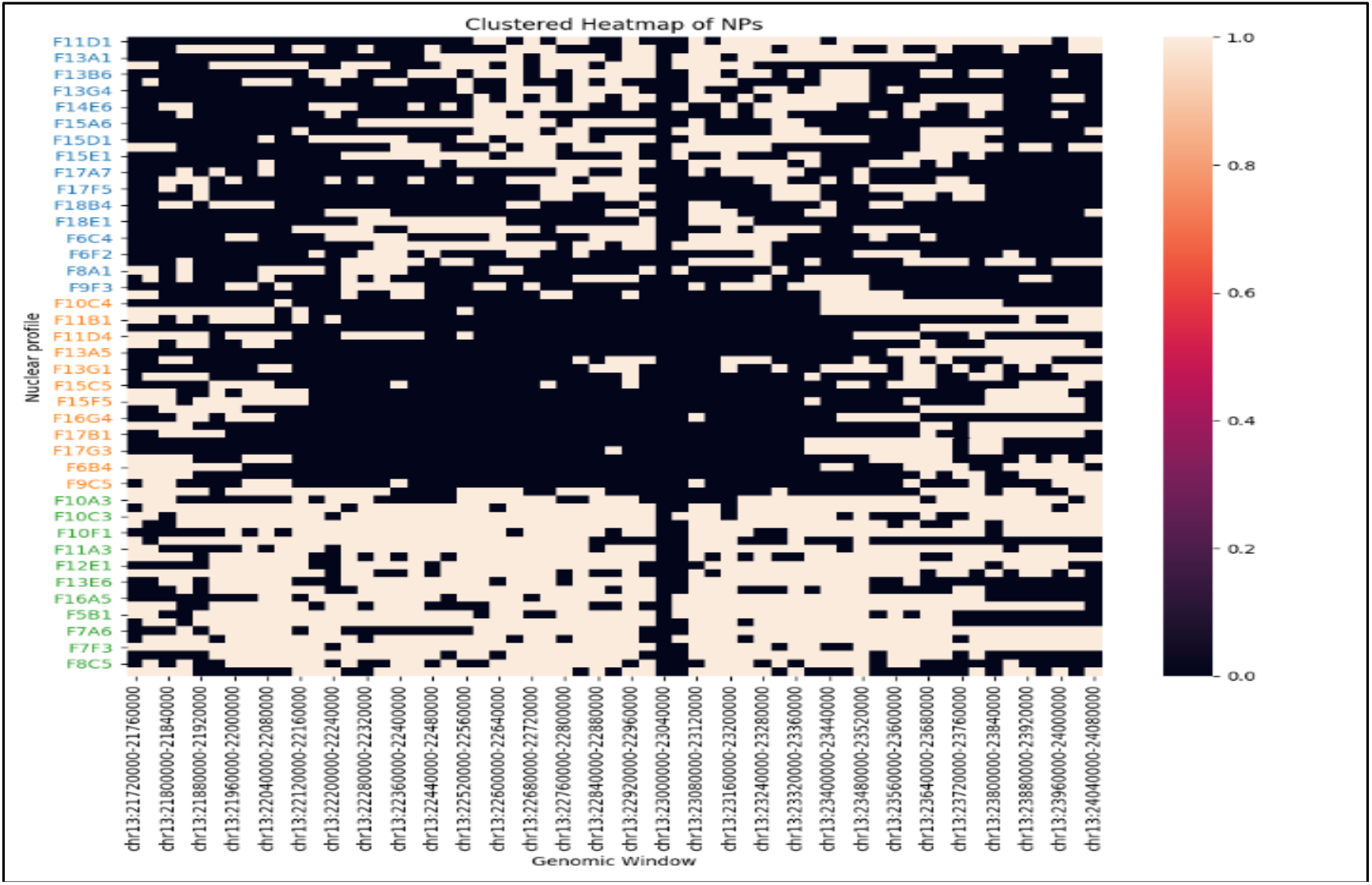
After clusters are obtained from k-means clustering, the segmentation table is grouped by cluster, with NP labels colored by the cluster in which they appear.

**Figure 5.**
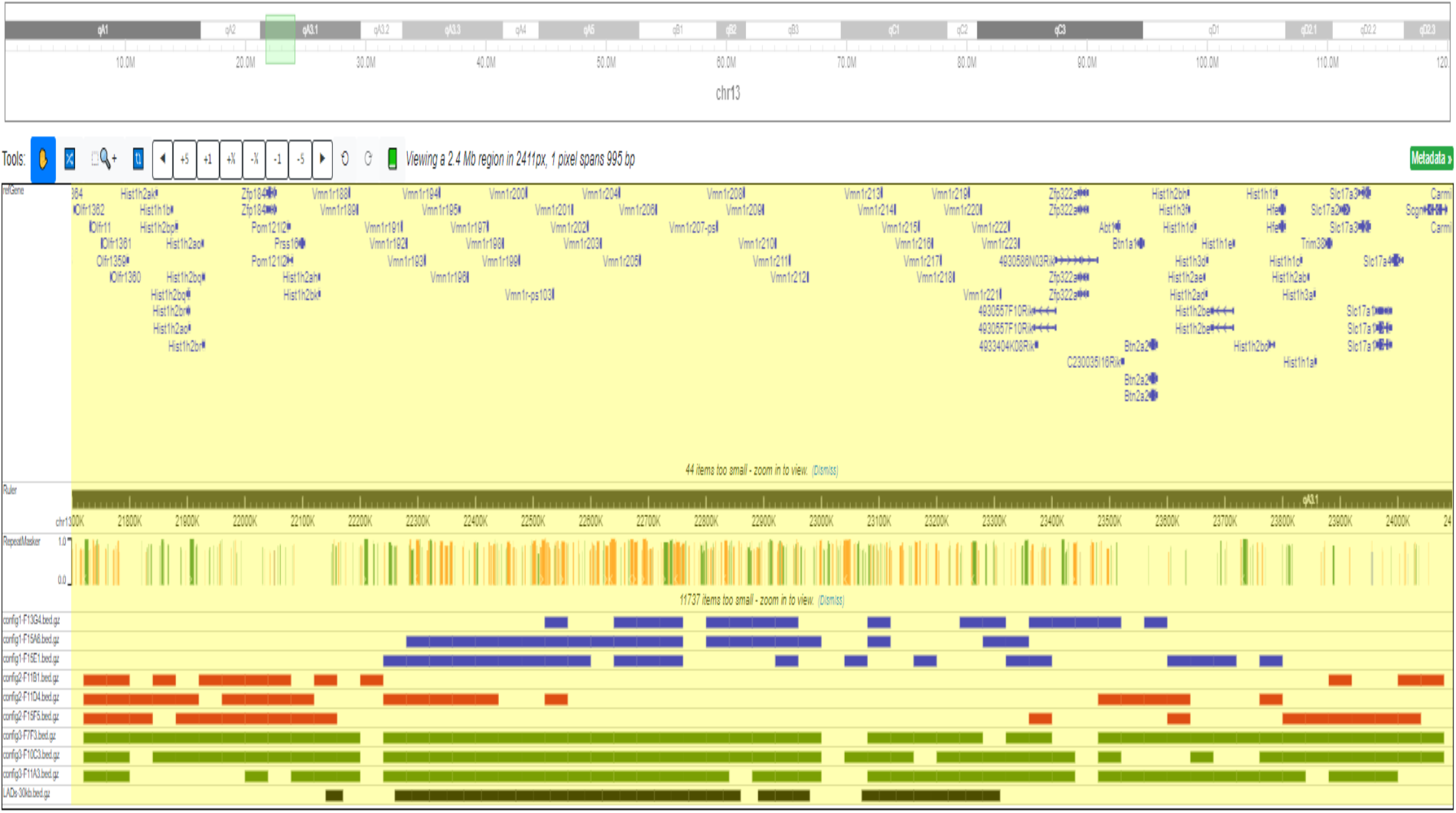
NPs shown as tracks in the Hist1 region. Three NPs from each cluster are depicted (cluster1=blue; cluster2=red; cluster 3=green). The black colored track indicates Lamina-Associated Domains (LADs).

We also explored whether the pseudo-circular shape in the PCA plot may represent the phases of the cell cycle by characterizing the activity of cell cycle genes [8]. For each cell cycle gene present in a window captured by an NP, the chromatin compaction of the genomic region containing the gene is estimated by counting the number of windows that the NP detected within the genomic neighborhood surrounding the cell-cycle-containing-window (we used 300Kb-sized neighborhoods and 1Mb-sized neighborhoods). This number is used as a proxy for compaction. Lower numbers indicate that the genes are in a decondensed genomic region, which is assumed to be in an **A** compartment and therefore active. A cell cycle is assigned to each NP by selecting the phase having genes with the highest de-compaction. The results (shown in Table 1) indicate that the largest group of cells in cluster 1 and cluster 2 contain active genes that are associated with the S phase. A strong indicator was not found for the phase captured by the cells in cluster 3.

**Table 1.** The numbers of de-compacted cell cycle loci in each cluster.

|  | **G1** | **S** | **G2** | **M** |
| --- | --- | --- | --- | --- |
| *Cluster 1* | 9 | 12 | 9 | 7 |
| *Cluster 2* | 6 | 11 | 8 | 3 |
| *Cluster 3* | 6 | 8 | 3 | 2 |

### 3.2 Feature generation

Since we are studying the HLB nuclear body, we examined an -omics feature that relates to nuclear organization - the Lamina Associated Domain (LAD). The presence of LADs is shown as a black track at the bottom of Figure 5. Additionally, the 3D slice position and chromatin compaction were estimated for each cluster.

The radial positions of all NPs were calculated in order to determine the distribution of the number of windows observed by NPs. Based on the distribution, the following cutoff points were determined: NPs that detected fewer than 2728 windows are considered to be apical; and NPs that detected more than 6057 windows are considered to be equatorial. Given these boundaries, the radial positions of the clusters can be estimated from the cluster-histograms shown in the Figure 6(a). Cluster 2 contains NPs that trend towards equatorial latitudes, and cluster 3 contains NPs that tend toward apical latitudes. The radial position of cluster 1 is more apical than cluster 2 and is less apical than cluster 3.

**Figure 6.**
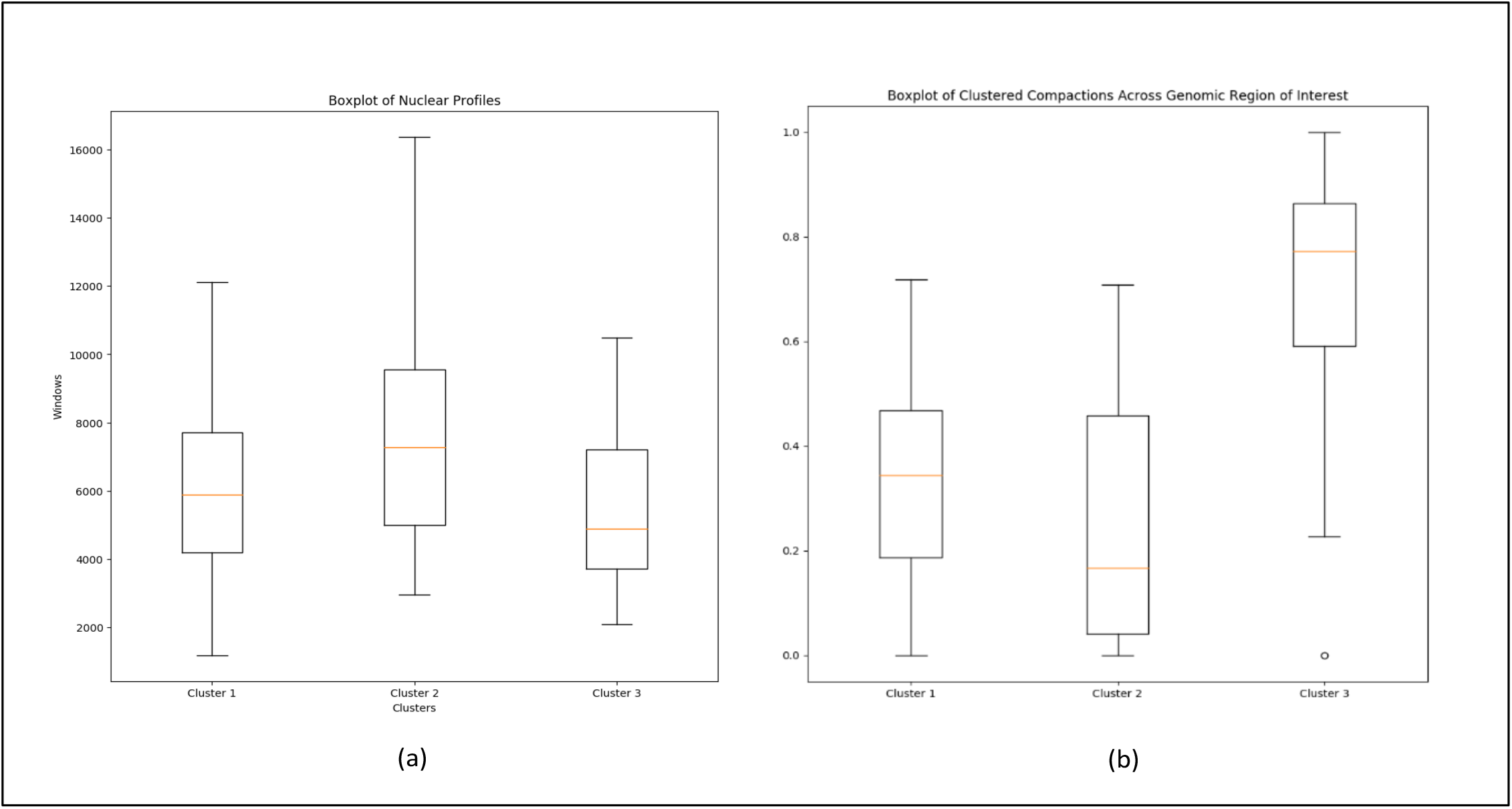
Characterizations of radial position and chromatin compaction of the NP clusters. (a) A summary of the number of windows in the NPs within each cluster. This number is used as a measure of radial position (e.g., equatorial NPs detect the largest numbers of windows). (b) Per cluster summaries of the window detection frequencies for the windows in the genomic region of interest.

Our geometric modeling approach was employed by plugging in the observed volume of a slice and the assumed parameters **a**,**b**,**c** (with a = 4, b=4, c=5.6), and solving for the values **n**,**m**,**k** to locate the position of a slice on the ellipsoid. We chose these values of a, b, and c because they are close to the dimensions of the sphere used in the original GAM model (diameter of 9 *μ*m) and this ellipsoid shape yields real-valued solutions for all of the observed slice volumes. However, we note that this choice of ellipsoid shape is a modeling assumption and impossible to rigorously determine from the slice data. We set **k** =1 and **m** =0.1 for all slices (analogous to choosing a particular rotation of the ellipsoid), and calculated **n**. As shown in Figure 7, the geometric model yields plausible locations for all nuclear profiles. All of the slices come from locations that capture a large part of the nucleus, consistent with the experimental procedure used to generate them. Additionally, we observe two distinct groups of slice orientations—those oriented along the major axis of the ellipsoid and those at an obtuse angle from the major axis.

**Figure 7.**
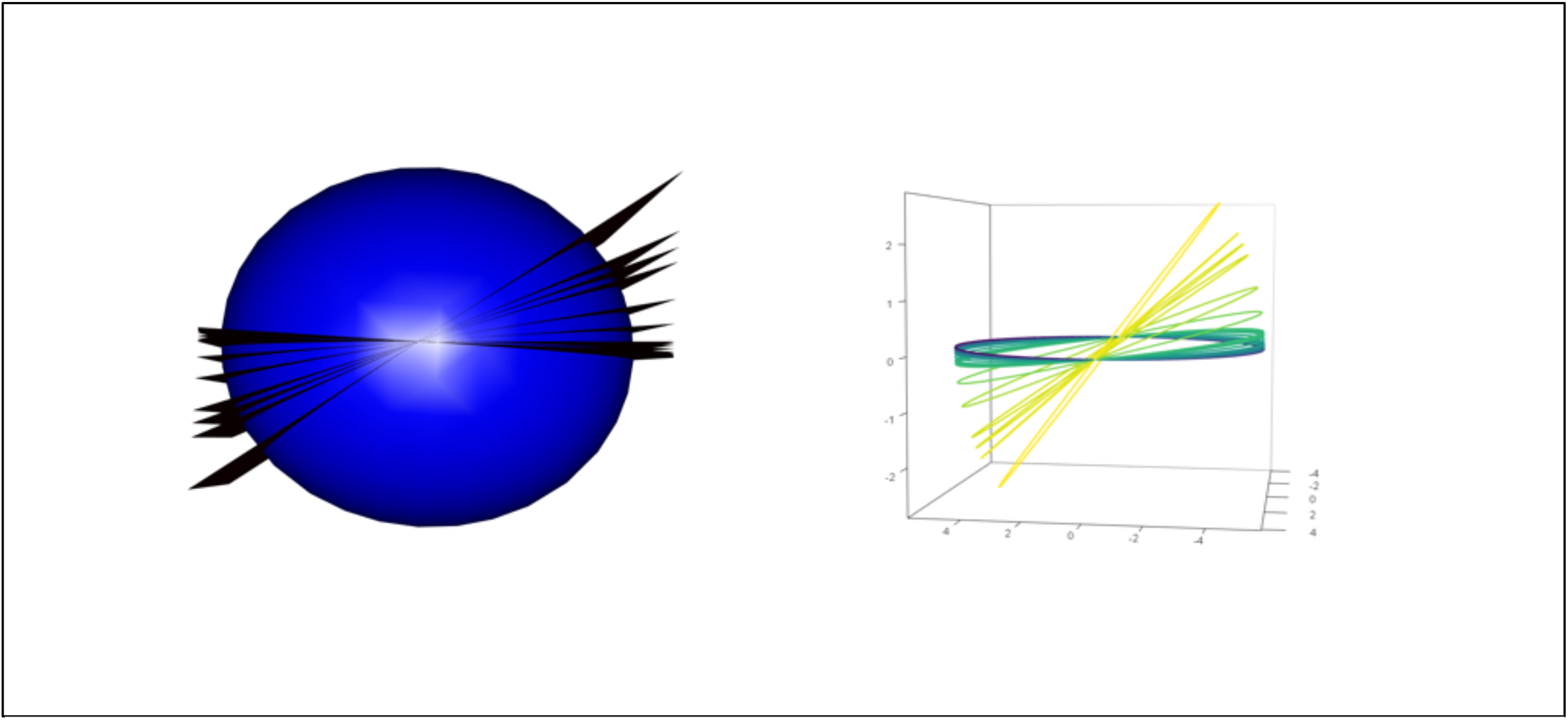
Plots of the 3D inferred slice locations using the ellipsoid model. Left: Each slice plotted as a plane. Right: The intersection curves between each slice and the ellipsoid, colored by the observed volume of the slice. Note that there are two distinct groups of slices, with different orientations and volumes.

To estimate the chromatin compaction of each cluster, **c**, we calculated the number of times that each 40Kb window was observed by the NPs *within* each cluster, **c** (i.e., we calculated the NP(**w**_**j,c**_) values, for all *j*; as described in section 2.2.2). The compaction estimates are shown in Figure 6(b), where it can be observed that the compaction of cluster 3 is high, and the compactions of cluster 1 and cluster 2 are low.

### 3.3 Interpretation

A strong, consistent biological narrative emerges from the *SluiceBox* analysis of the single-cell GAM model of the Histone Locus Body. In summary, the analysis reveals two cell states, each of which corresponds to a specific chromatin structure, radial position, and degree of compaction.

Figure 8 provides genome window interaction heatmaps for the NP clusters identified in the Hist1 locus. The chromatin interactions observed in each cluster are quite distinct. Observe the fluctuations in the structure of the HLB, with the regions in the middle of the Hist1 region interacting in panel *a* and the regions on the edges of the Hist1 region interacting in panel *b*. All regions interact in panel *c*. We found strong evidence that panels *a* and *b* capture one chromatin state, and that panel *c* captures a second chromatin state. Panel *d* combines the NPs from all three clusters, providing the view that pooled GAM would capture.

**Figure 8.**
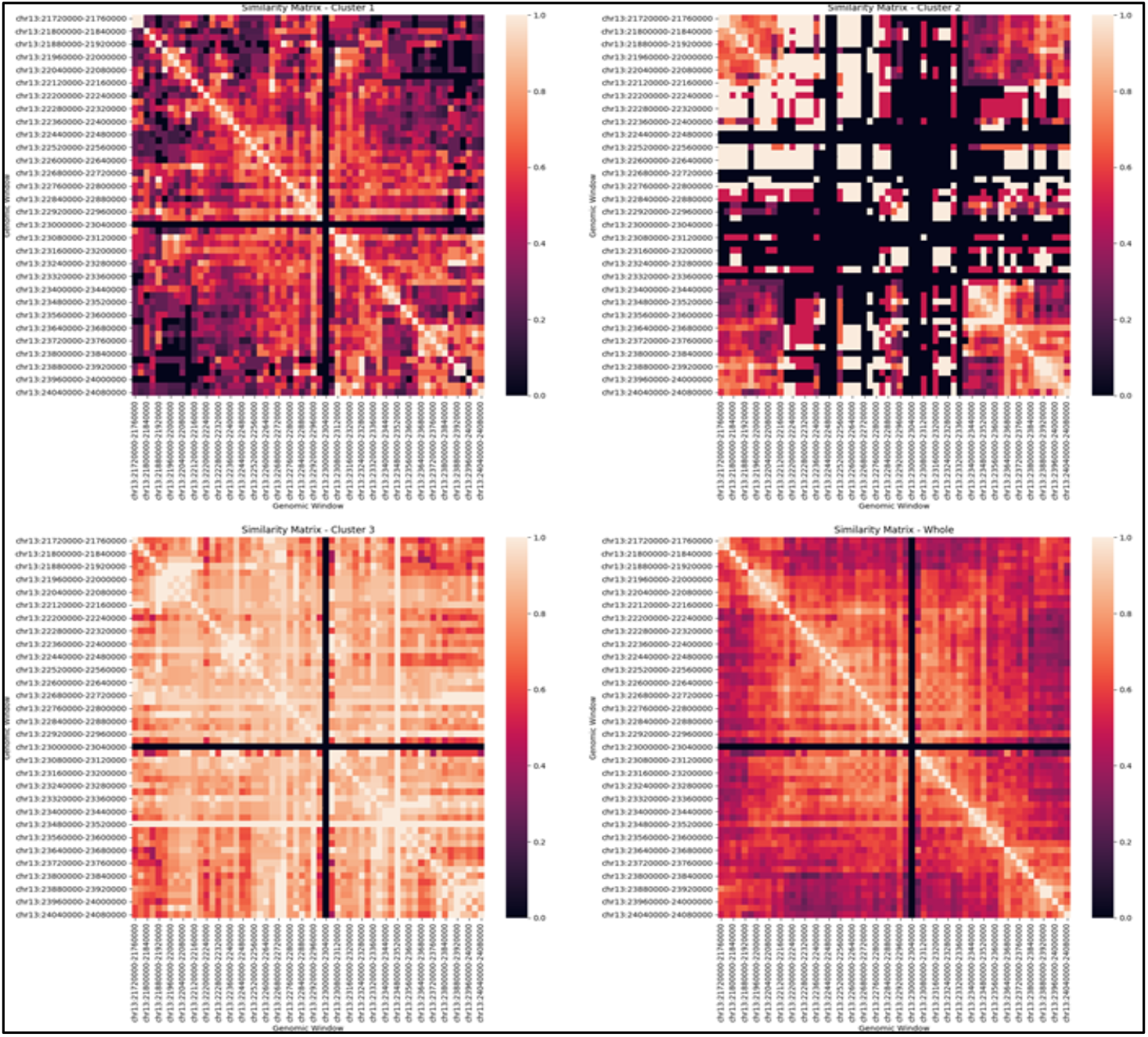
Different chromatin configurations detected by each cluster: cluster1 - panel (a); cluster2 - panel (b); cluster3 - panel (c); all clusters - panel(d).

The state captured by panels *a* and *b* depicts a loop, with panel *b* depicting the base of the loop (consisting of interacting histone gene groups from the two ends of the Hist1 region) and panel *a* depicting the closed portion of the loop (the lamina-associated portion of the region). Based on the radial position estimation, we can position the structure of the Hist1 loop within the nucleus (see Figure 9). The base of the loop (which contains the hist1 genes) is equatorial. The closed portion of the loop (which contains LADs) is in a peripheral region of the nucleus. The positioning and the lamina association are indicators that the function of the state represented by the loop between Hist1 gene groups that flank the LAD is to transcribe the hist1 genes. In contrast, the state captures in panel *c* can be interpreted as a state where all regions are condensed, whereby the Hist1 clusters are closer to the LAD (alternatively, this state might correspond to nuclear slices that fully contain the 2.5Mb region considered).

**Figure 9.**
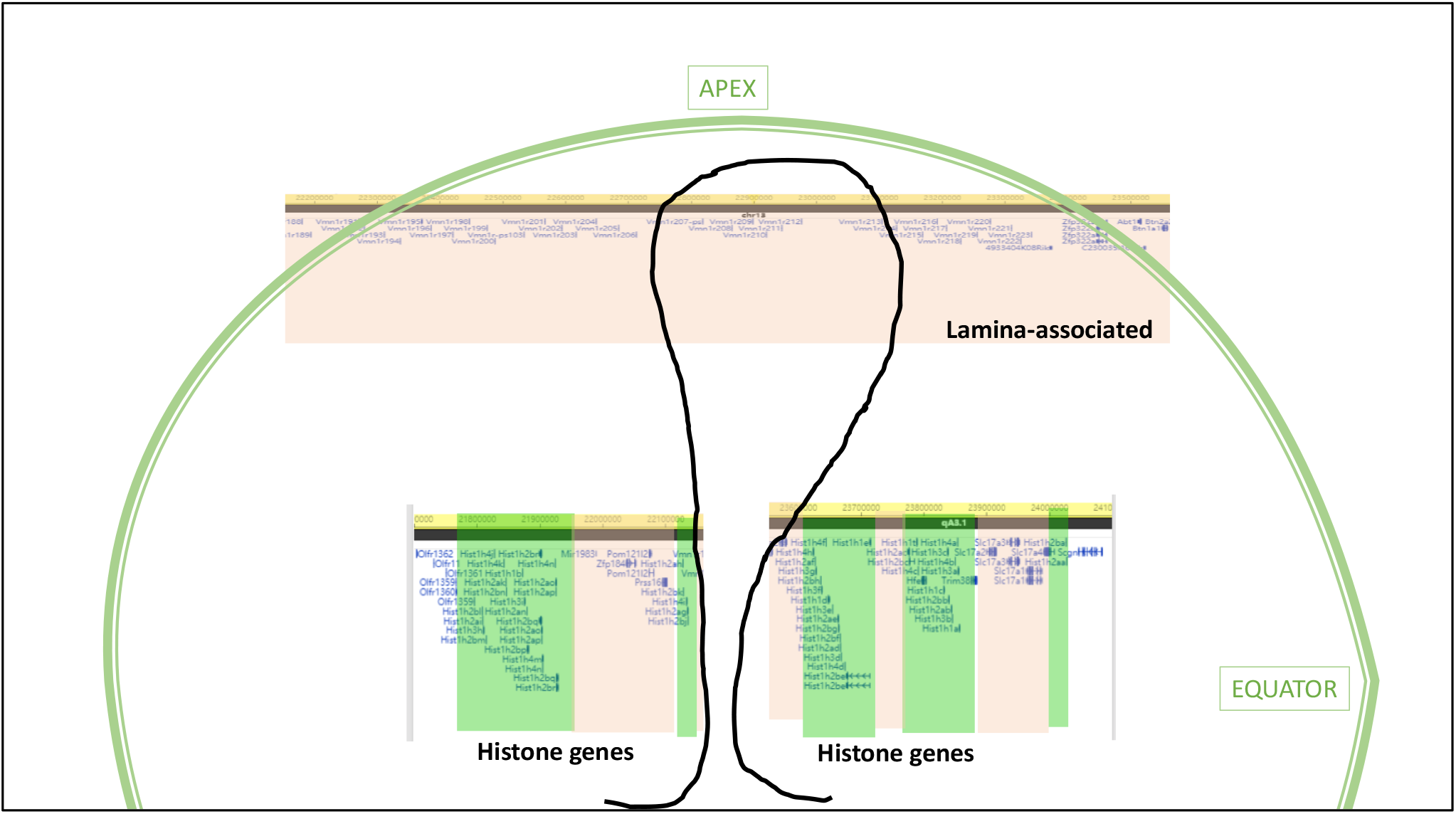
The structure inferred from clusters 1 and 2. Cluster 1 includes the center of the Hist1 region and is detected by slices that are more apical than cluster 2 slices. Experimental data indicate that the genomic windows in cluster 1 may also be lamina-associated. In cluster 2, regions containing the histone genes interact and are detected by de-condensed equatorial slices.

Consistent with this view, one can see that the chromatin is compacted in cluster 3 (Figure 6(b)) and that cluster 3 NPs tend to be apical (Figure 6(a)), as would be expected in chromatin that is associated with lamina in the nuclear periphery. Contrastingly, NPs in cluster 1 and cluster 2 contain de-condensed chromatin and tend to locate in latitudes of the nucleus that are more equatorial than the latitudes of NPs in cluster 3 (Figure 6). Cluster 2, which contains the histone genes, is more equatorial than cluster 1 – which is what would be expected when the histone genes (in cluster 2) are undergoing active transcription. The cell cycle analysis further confirms this interpretation, indicating that the NPs in clusters 1 and 2 are in S phase – the period of the cell cycle when histone genes are expressed.

## 4. Summary and Conclusions

Genome Architecture Mapping (GAM) has the ability to capture features that cannot be observed with ligation-based methods such as Hi-C. This paper presents *SluiceBox* a bioinformatics pipeline for performing single-cell analysis of GAM nuclear profile data. The utility of this approach is demonstrated by employing it to study an important genomic region, Hist1, which forms a Histone Locus Body. The process exploits single-cell information contained in GAM to study fluctuations in the chromatin structure of the HLB. Our results provide a demonstration and validation of the processes for gaining biological insights by identifying states of the HLB; discovering specific chromatin configurations of the HLB states; and identifying possible mechanism for formation of the HLB’s chromatin structure.

## Acknowledgements

LRW acknowledges support from the OHIO GERB Program. The work of WFM has been supported by NIH grant GM 29832, and the work of JDW has been supported by NIH grant R01 HG010883. AP acknowledges support from the Helmholtz Association (Germany) and the National Institutes of Health Common Fund 4D Nucleome Program grant U54DK107977.

## Notes

#### Summary of Updates

Added acknowledgements to sponsors.

